# Spatiotemporal characterization of corticolimbic dopamine and noradrenaline signaling and affective state modulation by mfb stimulation

**DOI:** 10.64898/2026.09.11.750608

**Authors:** Zhuo Duan, Yixin Tong, Andrada-Nicole Georgescu, Volker A. Coenen, Máté D. Döbrössy

## Abstract

**Background:** Deep brain stimulation of the medial forebrain bundle (mfb-DBS) is a promising intervention for psychiatric disorders, but the neurochemical mechanisms underlying its effects remain incompletely understood.

**Methods:** In adult Sprague–Dawley rats, fiber photometry was used to monitor mfb-DBS evoked dopamine (DA)- and noradrenaline (NA)- signaling in the nucleus accumbens (NAc) and prefrontal cortex (PFC), following different stimulation patterns, laterality conditions, and prolonged stimulation paradigms. Ultrasonic vocalizations (USVs) were recorded in parallel as a behavioral measure.

**Results:** Bilateral stimulation robustly increased catecholaminergic signals relative to sham in both NAc and PFC. The magnitude and temporal profile of these responses depended more on stimulation pattern rather than on laterality, with long pulse-width generally evoking the largest responses and laterality producing no significant differences across the four recording groups. During 10 min bilateral stimulation, catecholaminergic signals increased rapidly and remained elevated throughout stimulation, with the clearest sustained effect observed in NAc DA. mfb-DBS also increased USV calls and peak frequencies, whereas these behavioral changes showed no consistent region-specific correlation with catecholaminergic signal magnitude.

**Conclusions:** mfb-DBS recruits catecholaminergic signaling in a parameter-dependent and region-specific manner and increases positive-affective USV output. These findings identify temporal patterning as an important determinant of downstream circuit engagement, whereas the relationship between acute USV responses and regional catecholamine signaling appears more complex.

## Introduction

Direct electrical stimulation delivered through implanted electrodes has emerged as a promising approach for the treatment of severe neuropsychiatric disorders, particularly in patients who do not respond adequately to conventional therapies (Elias et al., 2021; Holtzheimer & Mayberg, 2011; Krauss et al., 2021; Sullivan et al., 2021). Among the targets explored for deep brain stimulation (DBS), the superolateral branch of the medial forebrain bundle (mfb), has attracted considerable interest because of its central position within reward- and motivation-related circuitry (Olds & Milner, 1954; Tong et al., 2023; Tozzi et al., 2024). Clinical studies in patients with treatment-resistant depression (TRD) (Coenen et al., 2019, 2022; Fenoy et al., 2018; Figee et al., 2022; Schlaepfer et al., 2013), together with preclinical work in rodents (Döbrössy et al., 2021; Furlanetti et al., 2015; Thiele et al., 2018; Tong et al., 2022), suggest that mfb-DBS can produce rapid and potentially sustained antidepressant effects. Despite these encouraging findings, the neurobiological mechanisms through which mfb-DBS influences affect-related circuits remain incompletely understood (Hariz et al., 2022).

One likely mechanism involves the recruitment of catecholaminergic signaling within mesocorticolimbic networks (Nutt, 2008; Robbins, 2018). Dopamine dysfunction has long been implicated in major depressive disorder, particularly in symptoms related to anhedonia, reduced motivation, and impaired reward processing (Grace, 2016). The nucleus accumbens (NAc) and prefrontal cortex (PFC) are key nodes within these circuits, and altered dopaminergic signaling in these regions has been linked to depressive phenotypes (Soares-Cunha et al., 2016; Robinson & Sohal, 2017; Höflich et al., 2019). Noradrenergic signaling also contributes to arousal (Poe et al., 2020), motivation, and affective state regulation (Rinaman, 2011), and growing evidence suggests that it may play an important role in antidepressant mechanisms (Schramm et al., 2001; Zhang et al., 2009). Together, these observations, supported by experimental data (Duan, Tong, et al., 2025; Duan, Zhao, et al., 2025; Miguel Telega et al., 2022), raise the possibility that mfb-DBS exerts part of its functional impact by engaging both dopamine- and noradrenaline-dependent signaling in the NAc and PFC.

A further unresolved issue is how stimulation parameters shape these neuromodulatory effects. The neural consequences of electrical stimulation depend not only on anatomical target location but also on stimulation pattern, temporal structure, and laterality (Ashouri Vajari et al., 2020; Grill, 2018; Mohan et al., 2020). Work in other neuromodulation settings, including theta-burst paradigms, has demonstrated that temporal patterning can strongly influence circuit responses (Huang et al., 2005; Lee et al., 2021; Solomon et al., 2021; Shaikh et al., 2024). However, it remains unclear how different mfb-DBS stimulation patterns recruit catecholaminergic signaling in downstream forebrain targets, whether these effects differ between unilateral and bilateral stimulation, and how they evolve during prolonged stimulation.

In the present study, we used fiber photometry to monitor dopamine- and noradrenaline- signals in the NAc and PFC during mfb-DBS under multiple stimulation conditions. We first determined whether bilateral mfb stimulation reliably recruits catecholaminergic responses relative to sham, and then compared distinct stimulation patterns and laterality conditions to assess how these parameters shape response magnitude and temporal dynamics. We further examined signaling during prolonged bilateral stimulation and explored the relationship between stimulation-evoked catecholaminergic responses and ultrasonic vocalization (USV) measures as an affect-related behavioral readout. By combining neurochemical and behavioral analyses, this study aims to clarify how mfb-DBS engages catecholaminergic circuits relevant to motivation and affect.

## Materials and Methods

### Animals

Sprague Dawley rats (SD, ♂: 16, ♀: 13, Charles River, Germany), aged 10-12 weeks and weighing 300–600 grams, were used in this study. Upon arrival at the facility, animals underwent a one-week acclimatization period. Rats were housed under a 12-hour light/dark cycle (lights on at 7:00 A.M.) with food and water available ad libitum. All procedures were conducted in accordance with the ARRIVE guidelines, approved by the local ethics committee, and complied with the ethical standards set by the Regierungspraesidium Freiburg (animal experiment application G21/144).

### Stereotaxic surgical preparation

Anesthesia was induced with 4 % isoflurane (2 l/min O_2_) in an induction box and maintained between 1.5 % and 2 % in a stereotactic frame (Stoelting, USA). Depth of anesthesia was monitored via the toe pinch reflex. Animals were randomly allocated and received GRAB sensors either AAV9- hsyn-NE2m (titer: 1.77 * 10^13^ particles/ml, YL003008, WZ Biosciences Inc., USA) or AAV9-hsyn- DA2m (titer: 1.77 * 10^13^ particles/ml, YL002009-AV9, WZ Biosciences Inc., USA) unilaterally in PFC (AP: + 3.0, ML: +0.4, DV:-3.2/-3.4, 500 nl each) or in NAC (site 1: 500 nl, AP: +1.6, ML: +1.1, DV: -6.8; site 2: 500 nl, AP:+1.2, ML: +1.1, DV: -6.9) via a 2 μl Hamilton syringe. The injections were controlled by a microsyringe pump (UMP3, World Precision Instruments) at 100 nl/min. Two weeks after the viral injection, bilateral DBS electrodes (Teflon coated, 90 % Platinum /10 % wire, 101-5T, Science Products) were implanted bilaterally into the mfb (AP: -2.8, ML: ±1.7, DV: -7.9). The optic fiber (400 μm, Doric Lenses, Canada) was positioned in the midpoint of the sensor virus injection site. Implants were secured with dental acrylic and bone cement (Palacos®, Heraeus Medical, Germany).

### Stimulation paradigm

After four weeks of viral sensor expression, each rat underwent titration to determine the optimal medial forebrain bundle deep brain stimulation (mfb-DBS) intensity. The appropriate stimulation current (ranging from 50 to 300 μA) was identified individually based on the expression of SEEKING behavior during stimulation, as previously described (Furlanetti et al., 2015). Each animal first underwent a 10-day intermittent stimulation phase, followed by a continuous stimulation phase (see Fig. 1). During the intermittent phase, rats received 5 seconds of mfb stimulation with a 50- second interstimulus interval, following an initial 50-second habituation period that served as baseline. This sequence was repeated 20 times per session. Post-stimulation recordings (50 seconds each) were collected immediately after the stimulation protocol. Three distinct stimulation patterns were tested: long pulse width DBS (LPW-DBS, 130 Hz and 100 μs), short pulse width DBS (SPW- DBS, 130 Hz and 50 μs), and intermittent theta burst DBS (iTB-DBS, 130 Hz and 100 μs with 3- pulse bursts at 5 Hz). For each stimulation pattern, DBS was delivered ipsilateral or contralateral to the fiber-photometry recording site—that is, in the mfb on the same or opposite hemisphere, respectively—or bilaterally to both mfb targets. All stimulation combinations were pseudorandomized and administered on consecutive days (see Fig. 1B). In vivo dopamine or noradrenaline release was recorded in the nucleus accumbens (NAc) and prefrontal cortex (PFC) during stimulation. Additionally, ultrasonic vocalizations were simultaneously recorded throughout the stimulation paradigm. After a one-week rest period following the intermittent stimulation phase, rats underwent chronic continuous stimulation. The protocol consisted of a single 50-second habituation period, which served as the baseline measurement, followed by two consecutive 10- minute sessions of continuous bilateral medial forebrain bundle stimulation (LPW-DBS). The two stimulation sessions were separated by a 60-second interval. Ultrasonic vocalizations were recorded continuously throughout the stimulation paradigm.

**Figure 1.**
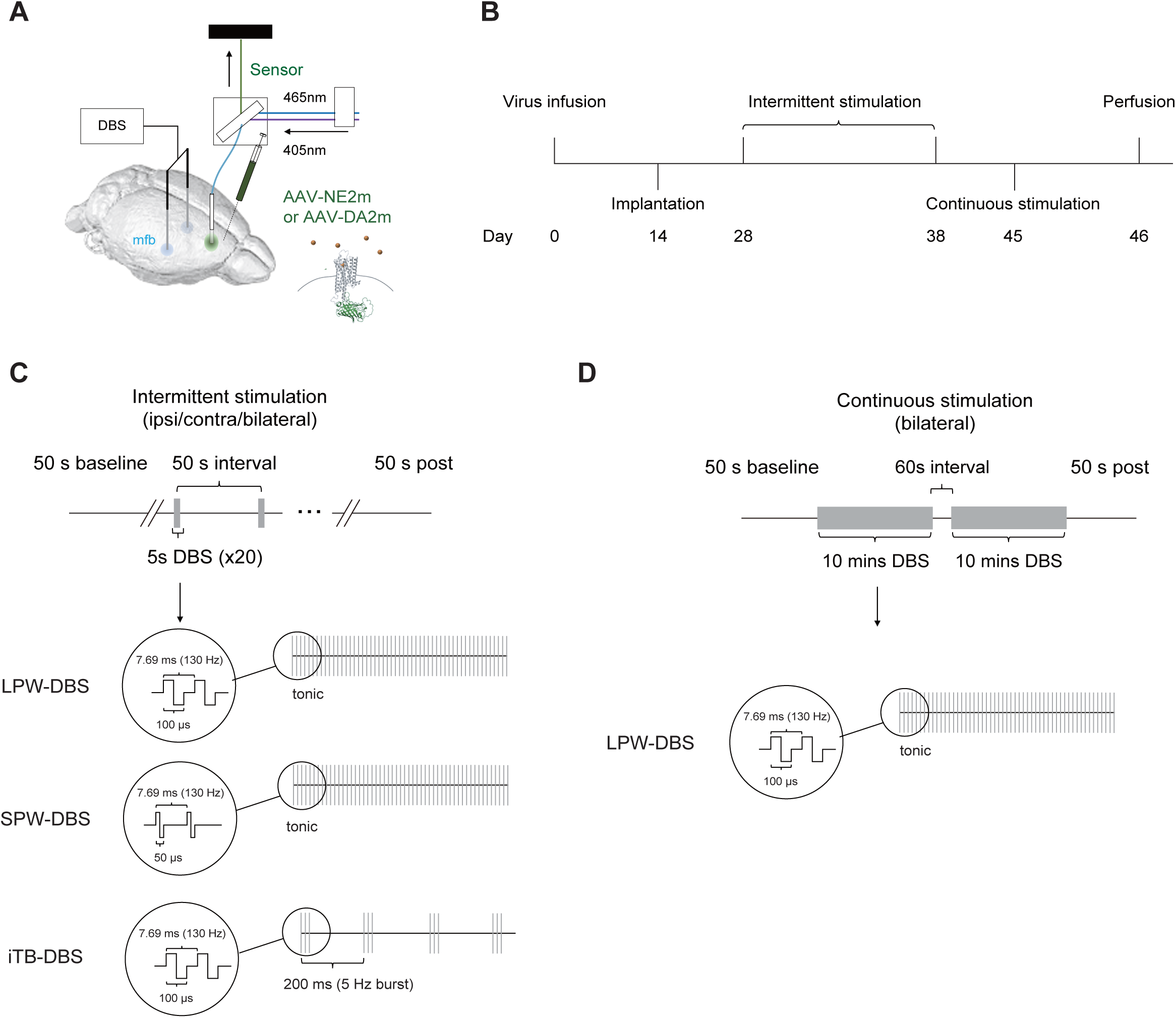
Overview of the fiber photometry and mfb-DBS experimental design. **(A)** Schematic of the recording and stimulation setup; **(B)** Experimental timeline showing viral infusion, implantation, intermittent stimulation experiments, continuous stimulation experiments, and perfusion; **(C)** Intermittent stimulation protocol. Trials consisted of a 50 s baseline, 5 s stimulation, 50 s interval, and 50 s post-stimulation epoch, repeated 20 times. Stimulation patterns included long pulse width DBS (LPW-DBS; tonic, 130 Hz, 100 μs), short pulse width DBS (SPW- DBS; tonic, 130 Hz, 50 μs), and intermittent theta-burst DBS (iTB-DBS; 3 pulses at 130 Hz per burst, repeated at 5 Hz, 100 μs pulse width). Intermittent stimulation was applied ipsilaterally, contralaterally, or bilaterally; **(D)** Continuous stimulation protocol. After a 50 s baseline, animals received two 10 min epochs of bilateral LPW-DBS separated by a 60 s interval, followed by a 50 s post-stimulation period. Continuous stimulation was delivered as tonic 130 Hz stimulation with a pulse width of 100 μs

### Fiber photometry recordings

Dopamine and noradrenaline signals were recorded during all stimulation paradigms. Prior to each session, all equipment was calibrated to ensure consistent LED power and detector sensitivity. Rats were connected to a fiber photometry patch cord and the DBS stimulator. The recording setup, as described elsewhere (Duan, Tong, et al., 2025; Miguel Telega et al., 2024), consisted of a fluorescence minicube, dual-wavelength LEDs with corresponding drivers, and a fluorescence detector (Doric Lenses® Inc., Canada). Photometric signals were excited at two wavelengths: 465 nm for GFP-sensitive excitation and 405 nm for isosbestic (GFP-insensitive) control. Data were acquired using the Synapse Suite software (Version 94, Tucker-Davis Technologies, USA) in conjunction with an RZ5 BioAmp processor. Custom-built stimulators generated transistor–transistor logic (TTL) pulses for DBS, which were synchronized with the photometric recordings via the RZ5 BioAmp system.

### Ultrasonic vocalization

Ultrasonic vocalizations (USVs) were recorded to assess the affective status of the rats. Recordings were acquired using an Ultramic UM192K microphone (Dodotronic, Italy) at a sampling rate of 0– 96 kHz. For analysis, the frequency band between 40 and 60 kHz—corresponding to positive affective vocalizations—was selected (Portfors, 2007; Wöhr & Schwarting, 2007). USV call number and call duration within this frequency range were detected using the DeepSqueak deep learning-based vocalization detector and subsequently validated manually (Coffey et al., 2019). Only animals classified as high callers, defined as those producing more than 20 detected calls, were included in the analysis (Schwarting et al., 2007).

### Histology

Following the final stimulation session, animals were terminally anesthetized by an overdose of 10% ketamine (Bela-Pharm GmbH & Co., KG, Germany) and 2% xylazine (Rompun, Bayer-Leverkusen, Germany) and intracardially perfused with ice-cold solution containing 4% paraformaldehyde (PFA) and 0.05% glutaraldehyde in 0.1 M phosphate buffered saline (PBS) at pH 7.4. The brains were removed from the skull, kept in 30% sucrose at 4°C until they sunk, and cut into 40 μm coronal sections. Viral expression and electrodes placement were verified by acquiring high-resolution images from ZEISS Axioscan 7 and analyzed using ZEN 3.8 software (Carl Zeiss).

### Statistical analysis

All data are presented as mean ± standard error of the mean (SEM) (Tables S1 and S2). Statistical analyses were performed using paired or unpaired Student’s t-tests, one-way ANOVA, one-way repeated-measures ANOVA, and Spearman’s rank correlation, as appropriate. Where applicable, repeated-measures ANOVA was followed by post hoc multiple-comparisons testing. Statistical analyses were conducted using Microsoft Excel and GraphPad Prism 9.0 (GraphPad Software). Figures were generated using GraphPad Prism 9.0 and MATLAB R2019a (MathWorks). Statistical significance was set at p < 0.05 and is indicated in the figures as P < 0.05, P < 0.01, P* < 0.001, and P < 0.0001.

## Results

### Bilateral mfb-DBS robustly increases catecholamine signaling relative to sham

To determine whether mfb-DBS reliably recruits catecholaminergic signaling, we first compared sham and bilateral LPW stimulation during fiber photometry recordings of dopamine (DA) and noradrenaline (NA) activity in the nucleus accumbens (NAc) and prefrontal cortex (PFC). Histological verification confirmed sensor expression and recording locations in NAc and PFC (Fig. 2A, D, G, J, Fig. S1). Across all four recordings, bilateral LPW stimulation evoked marked increases in z-scored fluorescence relative to sham, indicating robust stimulation-locked catecholamine responses in both accumbal and prefrontal sites (Fig. 2B, E, H, K).

**Figure 2.**
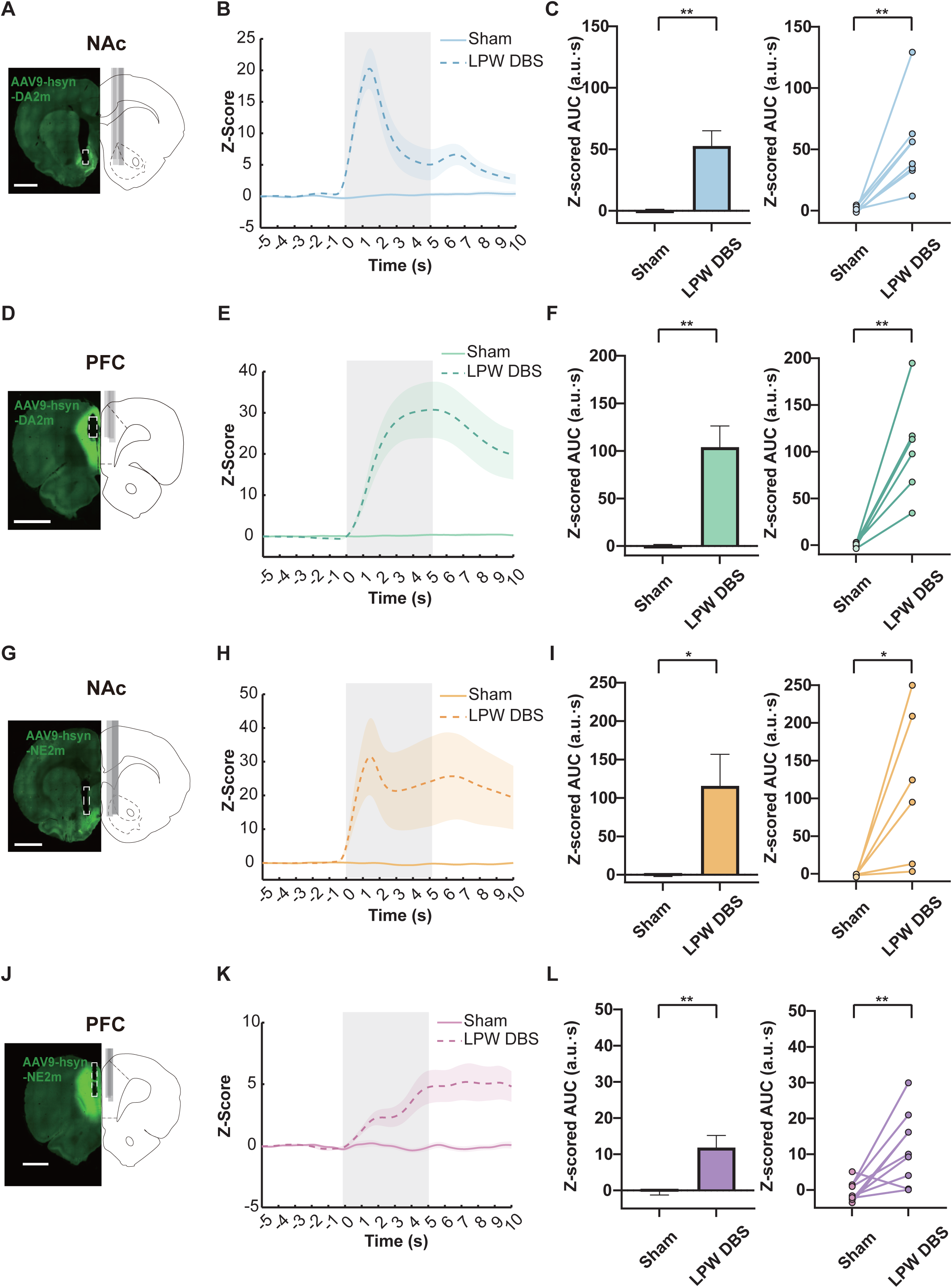
Bilateral long pulse width stimulation robustly increases catecholaminergic signaling in the NAc and PFC. **(A)** Representative histological verification of DA sensor expression and fiber placement in NAc. **(B)** Representative z-scored NAc DA fluorescence traces aligned to stimulation onset. Shaded area indicates the 5s stimulation period. **(C)** Stimulation-locked NAc DA z-scored area under the curve (AUC), shown as group summary and paired individual data. **(D)** Representative histological verification of DA sensor expression and fiber placement in PFC. **(E)** Representative z-scored PFC DA fluorescence traces aligned to stimulation onset. Shaded area indicates the 5s stimulation period. **(F)** Stimulation-locked PFC DA z-scored AUC, shown as group summary and paired individual data. **(G)** Representative histological verification of NA sensor expression and fiber placement in NAc. **(H)** Representative z-scored NAc NA fluorescence traces aligned to stimulation onset. Shaded area indicates the 5s stimulation period. **(I)** Stimulation-locked NAc NA z-scored AUC, shown as group summary and paired individual data. **(J)** Representative histological verification of NA sensor expression and fiber placement in PFC. **(K)** Representative z-scored PFC NA fluorescence traces aligned to stimulation onset. Shaded area indicates the 5s stimulation period. **(L)** Stimulation-locked PFC NA z-scored AUC, shown as group summary and paired individual data.

In NAc DA recordings, bilateral LPW stimulation produced a rapid and pronounced increase in signal relative to sham, characterized by a prominent early peak followed by a sustained elevation during the stimulation and early post-stimulation period (Fig. 2B). Quantification of the 5-s stimulation-locked response confirmed that z-scored AUC was significantly greater during LPW stimulation than during sham (Fig. 2C; paired t-test, t (7) = 4.211, p = 0.0040).

In PFC DA recordings, bilateral LPW stimulation also increased signal relative to sham, although the response profile appeared more sustained and of lower amplitude than that observed in NAc DA (Fig. 2E). Nevertheless, stimulation-locked z-scored AUC was significantly greater during LPW stimulation than during sham (Fig. 2F; paired t-test, t (5) = 4.771, p = 0.0050).

A robust LPW-evoked response was also observed in NAc NA recordings, in which stimulation induced a large increase in signal that remained elevated throughout the stimulation window and early post-stimulation period, whereas sham remained near baseline (Fig. 2H). This effect was confirmed by a significant increase in z-scored AUC during LPW stimulation relative to sham (Fig. 2I; paired t-test, t (5) = 2.846, p = 0.0360).

In PFC NA recordings, bilateral LPW stimulation produced a smaller but still clearly detectable increase in signal compared with sham (Fig. 2K). Quantification again demonstrated a significant increase in stimulation-locked z-scored AUC under LPW stimulation relative to sham (Fig. 2L; paired t-test, t (8) = 3.540, p = 0.0076).

Together, these findings establish that bilateral LPW-mfb-DBS robustly recruits both dopaminergic and noradrenergic signaling across NAc and PFC, thereby providing the foundation for subsequent analyses of how stimulation pattern and laterality shape these responses.

### Catecholamine responses during bilateral mfb-DBS depend strongly on stimulation pattern

We next examined how stimulation pattern influences catecholaminergic responses during bilateral mfb-DBS. Bilateral LPW, SPW, and iTB stimulation were compared during fiber photometry recordings of DA and NA activity in the NAc and PFC. Representative traces across recordings indicated that LPW generally elicited the strongest responses, whereas SPW and iTB produced smaller signals (Fig. 3A, E, I, M).

**Figure 3.**
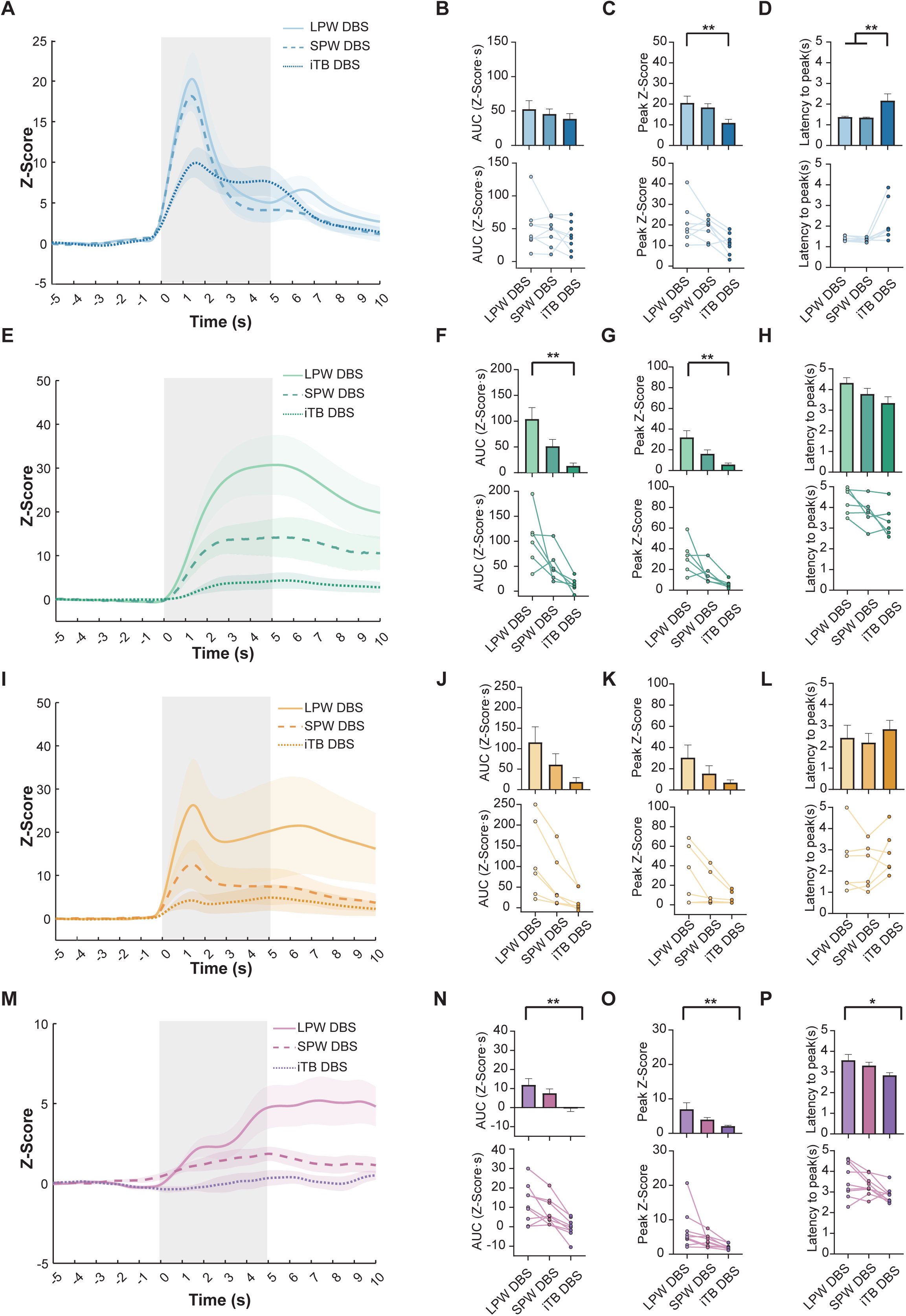
Comparison of stimulation patterns reveals distinct catecholaminergic dynamics in the NAc and PFC during mfb-DBS. **(A-D)** Data of NAc DA release in response to LPW-DBS, SPW-DBS, and iTB-DBS. **(A)** Representative z-scored fluorescence traces. **(B)** Stimulation-locked AUC. **(C)** Peak z-score. **(D)** Latency to peak response. **(E-H)** Data of PFC DA release in response to LPW-DBS, SPW-DBS, and iTB-DBS. **(E)** Representative z-scored fluorescence traces. **(F)** Stimulation-locked AUC. **(G)** Peak z-score. **(H)** Latency to peak response. **(I-L)** Data of NAc NA release in response to LPW-DBS, SPW-DBS, and iTB-DBS. **(I)** Representative z-scored fluorescence traces. **(J)** Stimulation-locked AUC. **(K)** Peak z-score. **(L)** Latency to peak response. **(M-P)** Data of PFC NA release in response to LPW-DBS, SPW-DBS, and iTB-DBS. **(M)** Representative z-scored fluorescence traces. **(N)** Stimulation-locked AUC. **(O)** Peak z-score. **(P)** Latency to peak response. The shaded region denotes the stimulation period. Statistical significance is indicated as shown.

In NAc DA recordings, bilateral LPW stimulation produced the largest response in the representative traces, whereas SPW and iTB evoked smaller signals (Fig. 3A). Despite this visual trend, z-scored AUC did not differ significantly across stimulation patterns (Fig. 3B; one-way ANOVA, F(2,21) = 0.5596, p = 0.5797). By contrast, stimulation pattern significantly affected peak amplitude (Fig. 3C; one-way ANOVA, F(2,21) = 4.385, p = 0.0256) as well as latency to peak (Fig. 3D; one-way ANOVA, F(2,21) = 5.652, p = 0.0109), with iTB reaching peak response later than LPW and SPW.

A similar but more pronounced pattern effect was observed in PFC DA recordings, in which LPW stimulation induced substantially larger responses than either SPW or iTB (Fig. 3E). This difference was reflected in a significant effect of stimulation pattern on both z-scored AUC (Fig. 3F; one-way ANOVA, F(2,15) = 8.889, p = 0.0028) and peak amplitude (Fig. 3G; one-way ANOVA, F(2,15) = 8.653, p = 0.0032). In contrast, latency to peak was less strongly affected and appeared comparatively similar across stimulation patterns (Fig. 3H; one-way ANOVA, F(2,15) = 3.023, p = 0.0789).

In NAc NA recordings, LPW again produced the largest response in the representative traces, while SPW and especially iTB evoked smaller signals (Fig. 3I). However, quantification did not reveal a significant effect of stimulation pattern on z-scored AUC (Fig. 3J; one-way ANOVA, F(2,15) = 3.051, p = 0.0773), peak amplitude (Fig. 3K; one-way ANOVA, F(2,15) = 2.058, p = 0.1622), or latency to peak (Fig. 3L; one-way ANOVA, F(2,15) = 0.4284, p = 0.6593). Thus, although descriptive trends were apparent, these differences did not reach statistical significance.

In PFC NA recordings, bilateral LPW stimulation also generated the largest responses, with SPW intermediate and iTB minimal responses (Fig. 3M). This pattern was confirmed by significant effects of stimulation pattern on both z-scored AUC (Fig. 3N; one-way ANOVA, F(2,24) = 6,270, p = 0.0064) and peak amplitude (Fig. 3O; one-way ANOVA, F(2,24) = 4.489, p = 0.0221). Latency to peak also differed across stimulation conditions in this recording, although these effects were smaller than those observed for response magnitude (Fig. 3P; one-way ANOVA, F(2,24) = 3.451, p = 0.0482).

Together, these findings indicate that catecholamine recruitment during bilateral mfb-DBS is modulated by stimulation pattern, with the effects varying across regions and response measures, including AUC, peak amplitude, and latency to peak. Pattern-dependent effects were most evident in PFC DA and PFC NA recordings, where LPW generally produced larger responses than SPW and iTB, whereas NAc recordings showed more limited or nonsignificant differences across stimulation conditions.

### mfb-DBS laterality does not significantly alter catecholaminergic responses in NAc or PFC

We next examined whether LPW-evoked catecholaminergic responses differ according to stimulation laterality. To address this, we compared bilateral, ipsilateral, and contralateral LPW stimulation during fiber photometry recordings of DA and NA activity in the NAc and PFC. Although representative traces showed some variation across conditions, these differences were not supported by statistical analysis in any of the four recording groups.

In NAc DA recordings, bilateral, ipsilateral, and contralateral LPW stimulation produced broadly similar response profiles in the representative traces (Fig. 4A). Consistent with this, z-scored AUC, peak amplitude, and latency to peak did not differ significantly across stimulation conditions (Fig. 4B–D).

**Figure 4.**
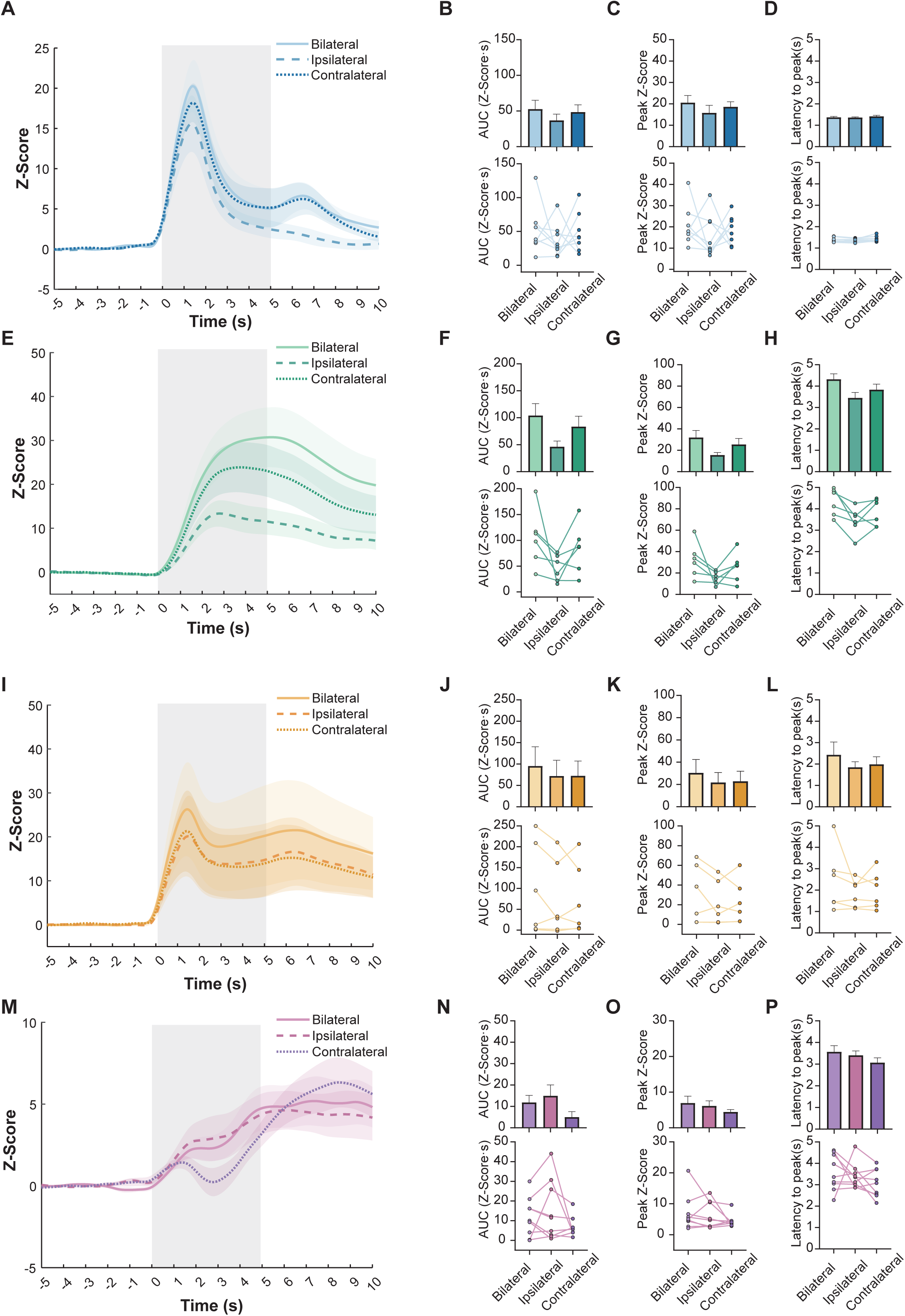
Laterality-dependent catecholaminergic responses to mfb-DBS in the NAc and PFC. **(A-D)** Data of NAc DA- responses in response to bilateral, ipsilateral, and contralateral stimulation. **(A)** Representative z-scored traces aligned to stimulation onset. **(B)** Stimulation-locked AUC. **(C)** Peak z-score. **(D)** Latency to peak response. **(E-H)** Data of PFC DA- responses in response to bilateral, ipsilateral, and contralateral stimulation. **(E)** Representative z-scored traces aligned to stimulation onset. **(F)** Stimulation-locked AUC. **(G)** Peak z-score. **(H)** Latency to peak response. **(I-L)** Data of NAc NA- responses in response to bilateral, ipsilateral, and contralateral stimulation. **(I)** Representative z-scored traces aligned to stimulation onset. **(J)** Stimulation-locked AUC. **(K)** Peak z- score. **(L)** Latency to peak response. **(M-P)** Data of PFC NA- responses in response to bilateral, ipsilateral, and contralateral stimulation. **(M)** Representative z-scored traces aligned to stimulation onset. **(N)** Stimulation-locked AUC. **(O)** Peak z-score. **(P)** Latency to peak response. The shaded region denotes the stimulation period. Statistical significance is indicated as shown.

Similarly, in PFC DA recordings, representative traces suggested some separation among bilateral, ipsilateral, and contralateral stimulation conditions, with bilateral stimulation appearing to evoke the largest response and ipsilateral stimulation the smallest (Fig. 4E). However, these differences were not supported by statistical analysis, as z-scored AUC, peak amplitude, and latency to peak did not differ significantly across laterality conditions (Fig. 4F–H).

In NAc NA recordings, bilateral, ipsilateral, and contralateral LPW stimulation again yielded partially overlapping response profiles, with only modest descriptive variation across conditions in the representative traces (Fig. 4I). Quantification likewise revealed no significant differences in z-scored AUC, peak amplitude, or latency to peak among the three stimulation conditions (Fig. 4J–L).

PFC NA recordings showed a similar pattern, with representative traces suggesting some visual differences across stimulation conditions but substantial overlap overall (Fig. 4M). As in the other recording groups, stimulation laterality did not significantly affect z-scored AUC, peak amplitude, or latency to peak (Fig. 4N–P).

Together, these findings indicate that, under LPW conditions, stimulation laterality does not significantly influence catecholaminergic recruitment in either NAc or PFC, in other words, unilateral and bilateral stimulation had comparable effects on the transmitter release.

### Catecholaminergic responses during prolonged bilateral LPW-DBS are most robust in NAc DA

To determine whether catecholaminergic responses were maintained during prolonged stimulation, we applied 10 min bilateral LPW-DBS and monitored DA and NA signals in the NAc and PFC. In all four recording groups, signal levels rose rapidly after stimulation onset and remained elevated throughout the stimulation period (Fig. 5A,C,E,G). In NAc DA recordings, repeated-measures analysis revealed a significant effect of time, and Dunnett’s post hoc test showed significant differences between the pre-DBS period and 1–10 min as well as the post-DBS period (Fig. 5B). In PFC DA, NAc NA, and PFC NA recordings, the traces showed a similar overall increase, but no individual time point differed significantly from pre-DBS after correction (Fig. 5D,F,H). Thus, the most consistent evidence for a sustained stimulation effect was observed in NAc DA.

**Figure 5.**
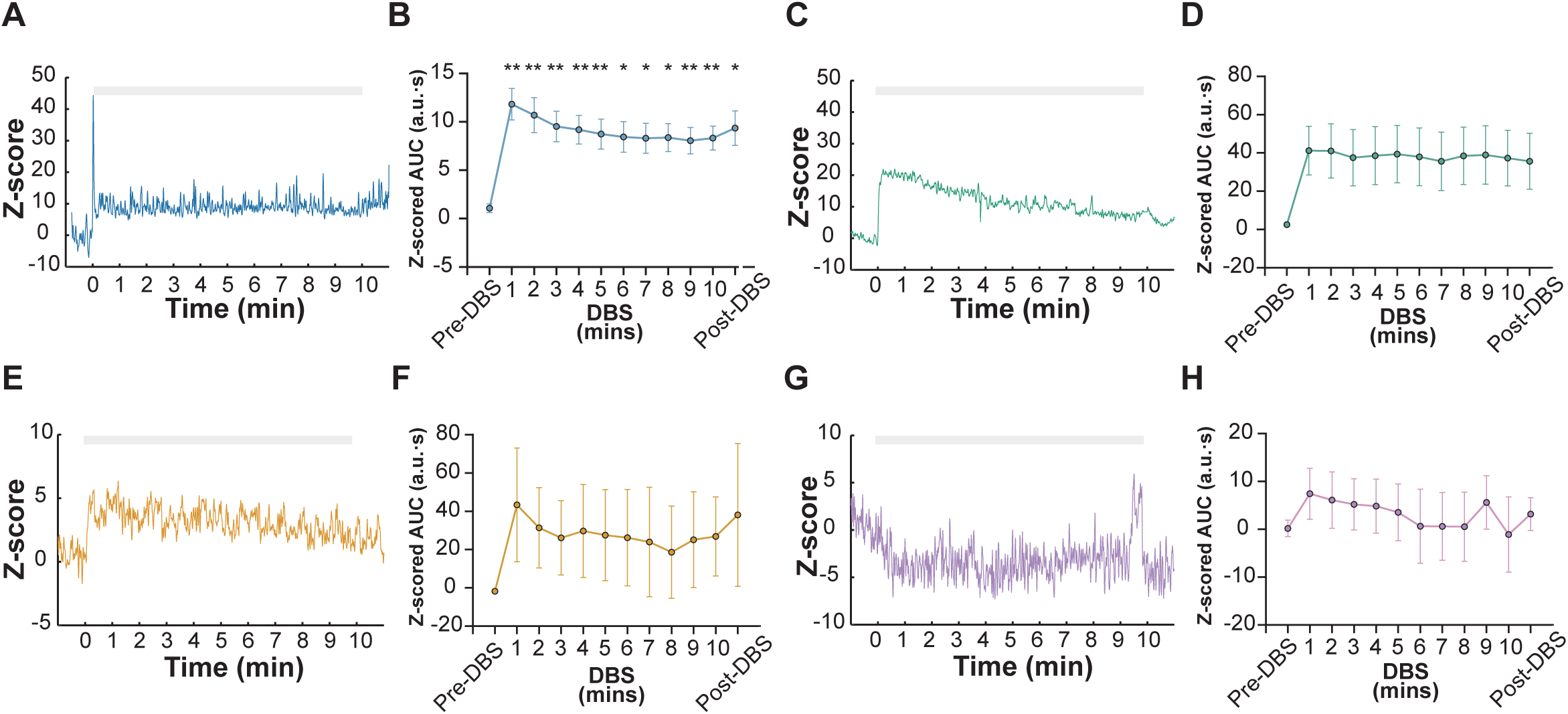
Catecholaminergic responses during prolonged bilateral mfb-DBS in the NAc and PFC. **(A-B)** NAc DA- responses during prolonged bilateral stimulation. **(A)** Representative z-scored fluorescence trace across the continuous stimulation session. **(B)** Minute-by-minute z-scored AUC before, during, and after DBS. **(C-D)** PFC DA- responses during prolonged bilateral stimulation. **(A)** Representative z-scored fluorescence trace across the continuous stimulation session. **(B)** Minute-by-minute z-scored AUC before, during, and after DBS. **(E-F)** NAc NA- responses during prolonged bilateral stimulation. **(E)** Representative z-scored fluorescence trace across the continuous stimulation session. **(F)** Minute-by-minute z-scored AUC before, during, and after DBS. **(G-H)** PFC NA- responses during prolonged bilateral stimulation. **(G)** Representative z-scored fluorescence trace across the continuous stimulation session. **(H)** Minute-by-minute z-scored AUC before, during, and after DBS. The shaded region denotes the stimulation period. Statistical significance is indicated as shown.

### Ultrasonic vocalizations and catecholamine signaling

To determine whether mfb-DBS alters affective vocal communication, we recorded ultrasonic vocalizations (USVs) during sham and DBS sessions across the different stimulation parameters (Figure 6A). DBS significantly increased the number of USV calls (Figure 6B; paired t-test, t(13) = 2.288, p = 0.0395) and USV peak frequency (Figure 6C; paired t-test, t(13) = 3.787, p = 0.0023) relative to sham, whereas USV call length showed no significant change (Figure 6D).

**Figure 6.**
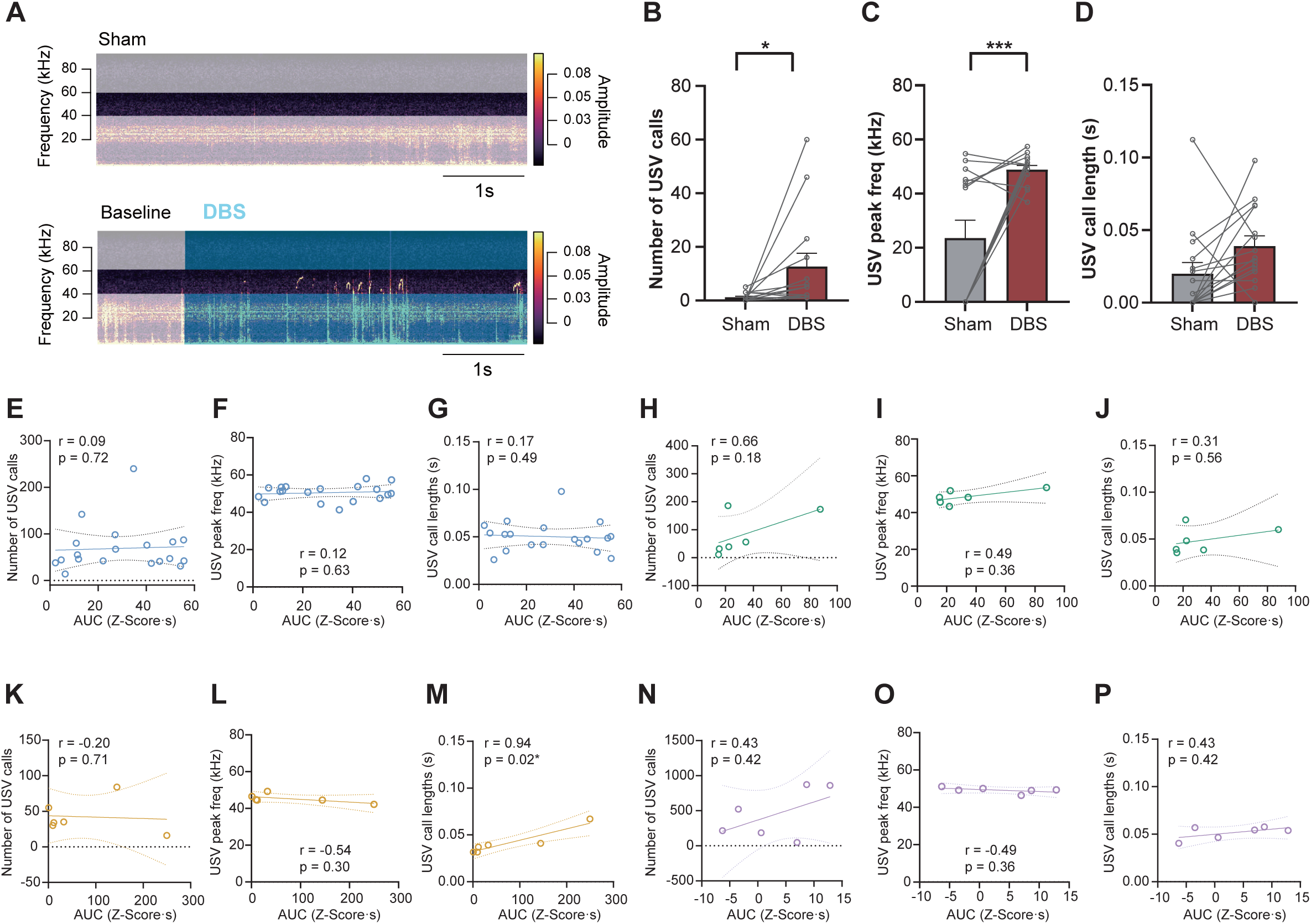
mfb-DBS increases ultrasonic vocalizations despite no region-specific correlation with catecholamine release. **(A)** Representative spectrograms of ultrasonic vocalizations (USVs) during Sham (top) and bilateral DBS (bottom) sessions, showing baseline (gray shading) and stimulation (teal shading) epochs. **(B– D)** Quantification of USV calls (B), peak frequencies (C), and call lengths (D) during Sham versus DBS sessions (paired comparisons; *p < 0.05, **p < 0.01, ***p < 0.001). **(E–G)** Correlations between NAc DA AUC (z-score·s) and USV calls (E), peak frequencies (F), and call lengths (G). **(H–J)** Correlations between PFC DA AUC and USV calls (H), peak frequencies (I), and call lengths (J). **(K–M)** Correlations between NAc NA AUC and USV calls (K), peak frequencies (L), and call lengths (M); **(N–P)** Correlations between PFC NA AUC and USV calls (N), peak frequencies (O), and call lengths (P). Dotted lines represent 95% confidence intervals of the linear regression fit; r and p values are shown for each Pearson correlation.

We next examined whether the magnitude of DBS-evoked transmitter release (AUC) was associated with these USV changes across the four catecholamine signals: NAc DA (Figure 6E–G), PFC DA (Figure 6H–J), NAc NA (Figure 6K–M), and PFC NA (Figure 6N–P). NAc DA AUC was not significantly correlated with the number of USV calls, peak frequency, or call length (Figure 6E–G; all P > 0.05). Likewise, PFC DA AUC showed no significant correlation with any USV parameter (Figure 6H–J; all P > 0.05).

NAc NA AUC was not significantly correlated with the number of USV calls or peak frequency (Figure 6K, L; both P > 0.05), but was significantly positively correlated with USV call length (Figure 6M; r = 0.94, P = 0.02). PFC NA AUC showed no significant correlation with any of the three USV parameters (Figure 6N–P; all P > 0.05).

Taken together, although mfb-DBS robustly enhanced ultrasonic vocal output, the magnitude of evoked catecholamine release was largely uncoupled from USV changes, with the exception of a selective association between NAc NA release and call length.

## Discussion

In this study, we showed that mfb-DBS recruits catecholaminergic signaling in the NAc and PFC in a stimulation-parameter dependent manner, with pattern exerting a stronger effect than laterality and prolonged bilateral stimulation producing a sustained response. Bilateral LPW stimulation reliably increased DA- and NA- signals relative to sham, whereas laterality effects under LPW conditions were not statistically significant. In contrast, stimulation pattern had clearer effects on response magnitude and timing, although these effects varied across brain regions and response measures. During prolonged bilateral LPW stimulation, all four recording groups showed sustained elevations in signal, with the clearest increase in NAc DA. Alongside these neurochemical changes, mfb-DBS also robustly enhanced ultrasonic vocalization, though this effect was largely dissociated from the magnitude of catecholamine release.

Three main observations stand out from these findings. First, the simultaneous recruitment of DA- and NA- signals in both the NAc and PFC supports the view that mfb-DBS acts on a distributed catecholaminergic network rather than on a single downstream target. This is consistent with the anatomical organization of the mfb as a heterogeneous fiber pathway linking midbrain structures with forebrain regions involved in reward, motivation, and affect (Coenen et al., 2012; Fenoy et al., 2022). From this perspective, the therapeutic effects reported with mfb-DBS may arise not from modulation of one anatomically discrete node, but from coordinated changes across several interconnected catecholaminergic targets. Such distributed engagement could be particularly relevant to psychiatric symptom domains, which are unlikely to reflect dysfunction in a single circuit element (Gálvez et al., 2015; Bewernick et al., 2017; Coenen et al., 2018; Remore et al., 2024).

Second, the stronger influence of stimulation pattern than laterality suggests that the physiological effect of mfb-DBS is determined less by the nominal side of stimulation than by the temporal features of the delivered current. This distinction is important because the mfb contains fibers with different excitability profiles, trajectories, and terminal fields. Altering pulse width or temporal organization may therefore change the subset of fibers recruited, the synchrony with which they are activated, or the extent to which downstream terminals are driven. The lack of clear laterality effects may reflect bilateral propagation of mfb-DBS effects through interconnected midbrain–forebrain circuits or convergence of ipsilateral and contralateral inputs onto common downstream targets. In support of this possibility, unilateral mfb-DBS has been reported to induce bilateral changes in gene-expression markers within frontal and striatal regions, indicating that stimulation effects can extend beyond the hemisphere containing the active electrode (Duan, Zhao, et al., 2025). It does not imply that laterality is unimportant in all settings, but indicates that temporal stimulation design may be the more tractable lever for shaping catecholaminergic output within the parameter range tested here. This interpretation aligns with evidence that waveform and temporal structure can modify neural responses even when stimulation location remains fixed (Grill, 2018; Ashouri Vajari et al., 2020; Gilbert et al., 2023).

Third, the regional heterogeneity observed across the NAc and PFC indicates that downstream targets do not simply mirror the activity of the stimulated tract. Instead, the final catecholaminergic response is likely shaped by target-specific processes, including local terminal density, uptake and autoregulatory mechanisms, receptor-mediated feedback, and the integration of converging inputs. The comparatively robust sustained NAc DA response may therefore identify accumbal dopamine as a particularly stable output of continuous mfb recruitment, whereas the more variable responses in the PFC and NA recordings may reflect stronger local regulation or greater dependence on the precise stimulation pattern. This distinction is biologically plausible given the different roles of the NAc and PFC in reward valuation, motivation, cognitive control, and affect regulation (Piantadosi et al., 2020; Friedman & Robbins, 2022). More broadly, it suggests that mfb-DBS should not be understood as producing a single “catecholamine effect,” but as shaping a set of regionally differentiated neurochemical states.

The heterogeneous pattern of catecholaminergic recruitment observed in this study may in part reflect the fiber composition of the mfb itself. The mfb is not a uniform tract but a compact bundle of ascending and descending axons belonging to multiple neurotransmitter systems, including dopaminergic, noradrenergic, and serotonergic fibers, as well as fibers of passage unrelated to catecholaminergic transmission (Coenen et al., 2012; Veening et al., 1982). These axonal populations differ in diameter, myelination, trajectory, and proximity to the electrode. Catecholaminergic fibers traversing the mfb include small-diameter and unmyelinated axons, whereas larger myelinated fibers of passage are also present within the tract (Nirenberg et al., 1996; Yeomans, 1989). Because axon diameter and myelination strongly influence excitability during extracellular stimulation, larger myelinated axons are generally activated at lower thresholds than thinner or unmyelinated fibers (Grill, 2018; McIntyre & Grill, 2002). The distinct DA- and NA- signals observed across the NAc and PFC may therefore reflect selective engagement of fiber populations within the stimulated volume, rather than equivalent activation of all mfb-traversing axons.

The stimulation-pattern effects observed here are consistent with this selective-recruitment framework. Relative to shorter pulse widths, longer pulses can facilitate activation of axons with longer membrane time constants or higher excitation thresholds by allowing more time for charge accumulation (Grill, 2018; Ranck, 1975). Conversely, SPW stimulation is expected to preferentially recruit larger, more excitable fibers closer to the electrode while sparing smaller axons that require longer charge delivery to reach threshold (McIntyre & Grill, 2002). The generally larger responses evoked by LPW are therefore compatible with broader recruitment of mfb elements that are less readily activated by SPW stimulation. This could include smaller-diameter catecholaminergic axons, although the present data do not identify the fibers responsible. The more pronounced LPW advantage in the PFC further suggests that pathways contributing to prefrontal DA- and NA- signals may be particularly sensitive to pulse-width-dependent recruitment, or that local prefrontal terminal mechanisms amplify differences in upstream drive. iTB provides a complementary example of how temporal structure may shape downstream output within the same anatomical target. Its brief high- frequency bursts and intervening silent periods differ from continuous LPW and SPW stimulation in the continuity and temporal organization of axonal and terminal activation. This type of theta-burst structure was originally designed to mimic endogenous theta-frequency activity and can engage short-term synaptic plasticity differently from continuous stimulation (Grill, 2018; Huang et al., 2005). The region-specific iTB effects observed here are compatible with altered synchrony of recruited fibers, incomplete buildup of sustained terminal drive, and/or changes in presynaptic release dynamics across bursts. In NAc DA, the dissociation between peak amplitude and AUC is consistent with a temporal redistribution of the response rather than a simple reduction in cumulative signal. By contrast, the more marked attenuation of PFC DA- and NA- responses suggests that circuits contributing to prefrontal catecholamine signaling may depend more strongly on continuous activation. These interpretations remain tentative, but they indicate that burst structure can influence not only the magnitude of catecholaminergic signaling, but also its temporal profile and regional expression.

The prolonged-stimulation findings suggest that continuous mfb-DBS does not simply maintain a uniform catecholaminergic state across downstream targets. Instead, the persistence of stimulation- evoked signaling appears to be shaped by target-specific mechanisms that regulate transmitter dynamics over time. The comparatively robust NAc DA response may reflect the predominant role of the dopamine transporter (DAT) in terminating dopaminergic signaling in this region, in contrast to the PFC, where DAT expression is low and dopamine clearance instead relies primarily on the norepinephrine transporter (NET) and catechol-O-methyltransferase (COMT) (Morón et al., 2002). These distinct clearance mechanisms could allow NAc DA signaling to be replenished and sustained more effectively during continuous mfb stimulation, whereas slower or less efficient PFC dopamine clearance may promote greater local feedback and stimulus-specific adaptation. Consistent with this possibility, repeated activation of prefrontal dopaminergic afferents produces adaptive, attenuated dopamine responses in the PFC that are not observed in the NAc (Jackson & Moghaddam, 2004).

This distinction is relevant to the proposed therapeutic actions of mfb-DBS because the NAc occupies a central position in motivational and reward-related processing; sustained engagement of accumbal DA signaling could therefore represent one mechanism through which continuous mfb stimulation influences motivational or affective states. However, the present recordings cover only 10 min and cannot determine whether this response is maintained during chronic DBS or whether it directly mediates antidepressant-like effects.

These findings also have implications for DBS parameter optimization. Clinical and preclinical DBS outcomes are known to be sensitive to stimulation parameters (Ramasubbu et al., 2018), but mechanistic comparisons across parameter sets remain limited for mfb stimulation (Ashouri Vajari et al., 2020; Kong et al., 2019; Miguel Telega et al., 2025). The present data suggest that temporal patterning is an important determinant of catecholaminergic recruitment, whereas laterality appears to play a more limited role. Because temporal patterning, rather than laterality, is the primary driver of this recruitment, this asymmetry in parameter sensitivity may help explain why stimulation protocols targeting the same anatomical pathway can nonetheless produce divergent physiological effects. More broadly, the results argue against viewing mfb-DBS as a unitary intervention and instead support a framework in which stimulation outcome reflects the interaction between target anatomy and stimulation design, a distinction with direct relevance for individualized parameter selection in clinical practice.

Beyond its neurochemical effects, mfb-DBS also produced a clear behavioral signature: DBS significantly increased the number of USV calls and USV peak frequency relative to sham, indicating that stimulation of this pathway reliably enhances affective vocal output. However, when we examined whether the magnitude of evoked catecholamine release was associated with the magnitude of these behavioral changes, we found that USV parameters were largely uncoupled from transmitter AUC across all four recording groups, with the exception of a selective positive correlation between NAc NA release and USV call length. This dissociation suggests that the occurrence of a behavioral response and the magnitude of that response may be governed by partially distinct mechanisms: we speculate that the presence of USV changes may depend on whether a given stimulation pattern engages the relevant circuit above some functional threshold, whereas the size of the neurochemical signal itself may be a comparatively poor predictor of downstream behavioral output once that threshold is crossed. The selective association observed for NAc NA and call length raises the possibility that noradrenergic signaling in the NAc contributes specifically to the temporal structuring of vocal output, a function distinct from simply triggering vocalization. This interpretation is consistent with pharmacological evidence that noradrenergic receptor manipulation selectively alters USV duration independent of call rate (Grant et al., 2018), although the present interpretation remains speculative given the limited sample size for this comparison.

Several limitations should be considered. First, the sample sizes were modest, which likely reduced power for some comparisons, particularly in the laterality and prolonged-stimulation analyses. Second, the present study focused on acute and subacute photometry responses and therefore does not address longer-term adaptations to repeated or chronic stimulation. Third, USV recordings were obtained under acute stimulation conditions and from a subset of animals in which vocalization occurred reliably enough to permit correlation analyses; several stimulation-pattern and animal combinations yielded few or no calls, which limited statistical power for detecting AUC–behavior associations within individual groups and may have contributed to the largely null correlation findings.

Taken together, these findings indicate that mfb-DBS does not produce a uniform catecholaminergic response, but instead recruits DA and NA signaling in a manner that depends on stimulation pattern, brain region, and stimulation duration. Across the conditions tested in this study, stimulation pattern had a clearer impact than laterality, and prolonged stimulation showed the most robust sustained effect in NAc DA. In parallel, mfb-DBS also significantly increased ultrasonic vocal output, but this behavioral effect was largely dissociated from the magnitude of evoked catecholamine release. These results provide a mechanistic basis for refining mfb-DBS protocols and support the idea that temporal design is a critical determinant of downstream circuit engagement, comparable in importance to the choice of stimulation laterality.

## Acknowledgments

We would like to thank Johanna Wessolleck and Jasmin Weis (lab of SIN, University of Freiburg Medical Center, Germany) for their technical support; Gerd Strohmeier (Werkstatt, University of Freiburg Medical Center, Germany) for providing and helping out with the DBS hardware; Dr. Ute Häußler, Experimental Epilepsy Research Group and Lighthouse Core Facility (University of Freiburg Medical Center, Germany) for sharing their microscopy facilities. We thank Yuzehngheng Zhang for helping visualization. Open Access funding enabled and organized by Projekt DEAL.

## Author contributions

Z.D. and Y.T. performed the experiment, analyzed the data and generated figures; A.G assisted the data analysis. Z.D. and Y.T. wrote the first version of the manuscript and with M.D., and V.A.C. synthesized the final manuscript. All the authors read and agreed with the manuscript.

## Data and code availability

Data reported in this paper will be shared by the lead contact upon request. Any additional information required to reanalyze the data reported in this paper is available from the lead contact upon request.

## Funding and Discloser

This work was supported by the Department of Stereotactic and Functional Neurosurgery, Medical Center – University of Freiburg, Germany, and by the UPSIDE project, funded by the European Union’s Horizon Europe EIC Pathfinder programme under grant agreement No. 101070931. Y.T. received support from the Hans A. Krebs Medical Scientist Program and Christian Homann Stiftung.

