## Supplementary figures and images for "Spatiotemporal characterization of corticolimbic dopamine and noradrenaline signaling and affective state modulation by mfb stimulation"

### Supplemental Figure 1

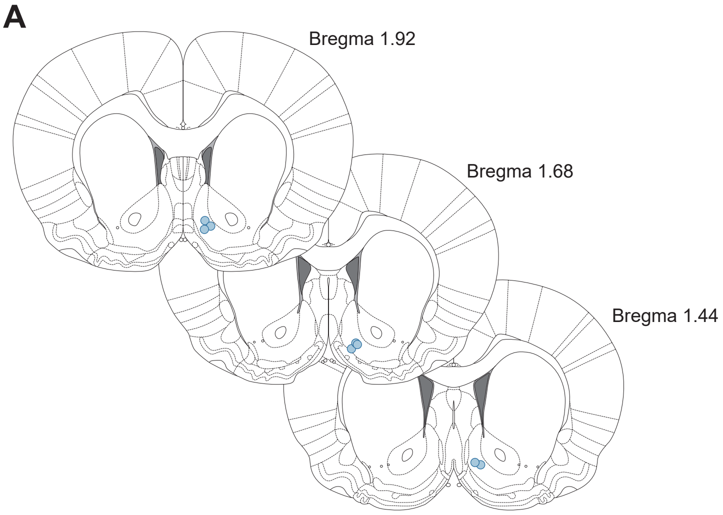

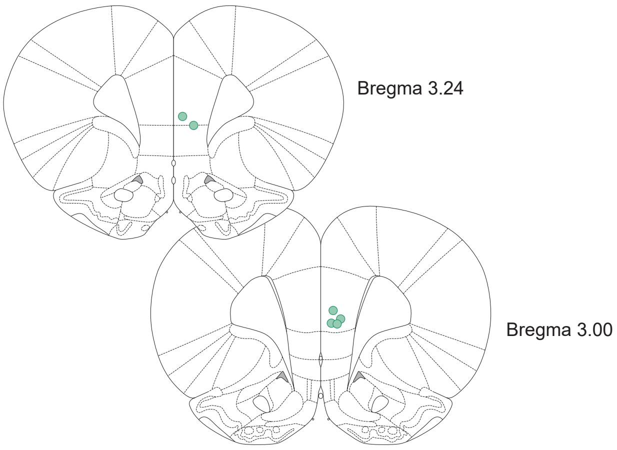

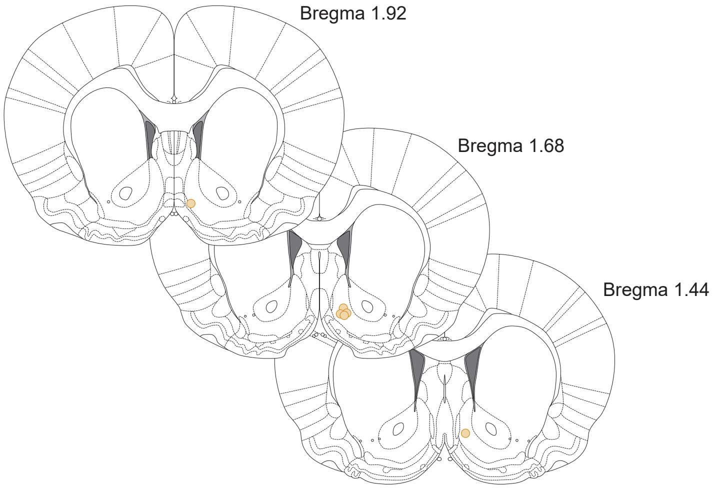

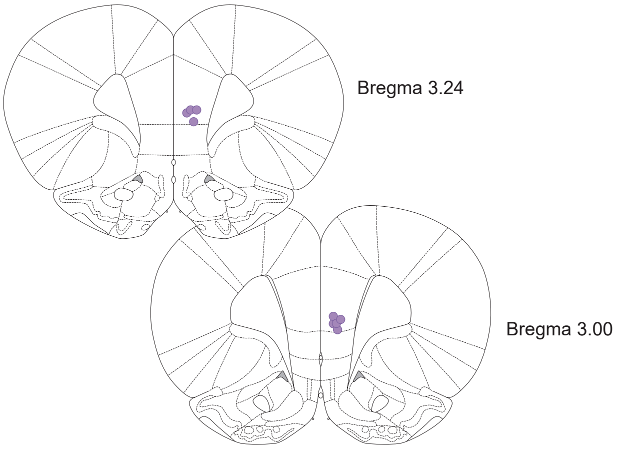

### Supplemental Figure 2

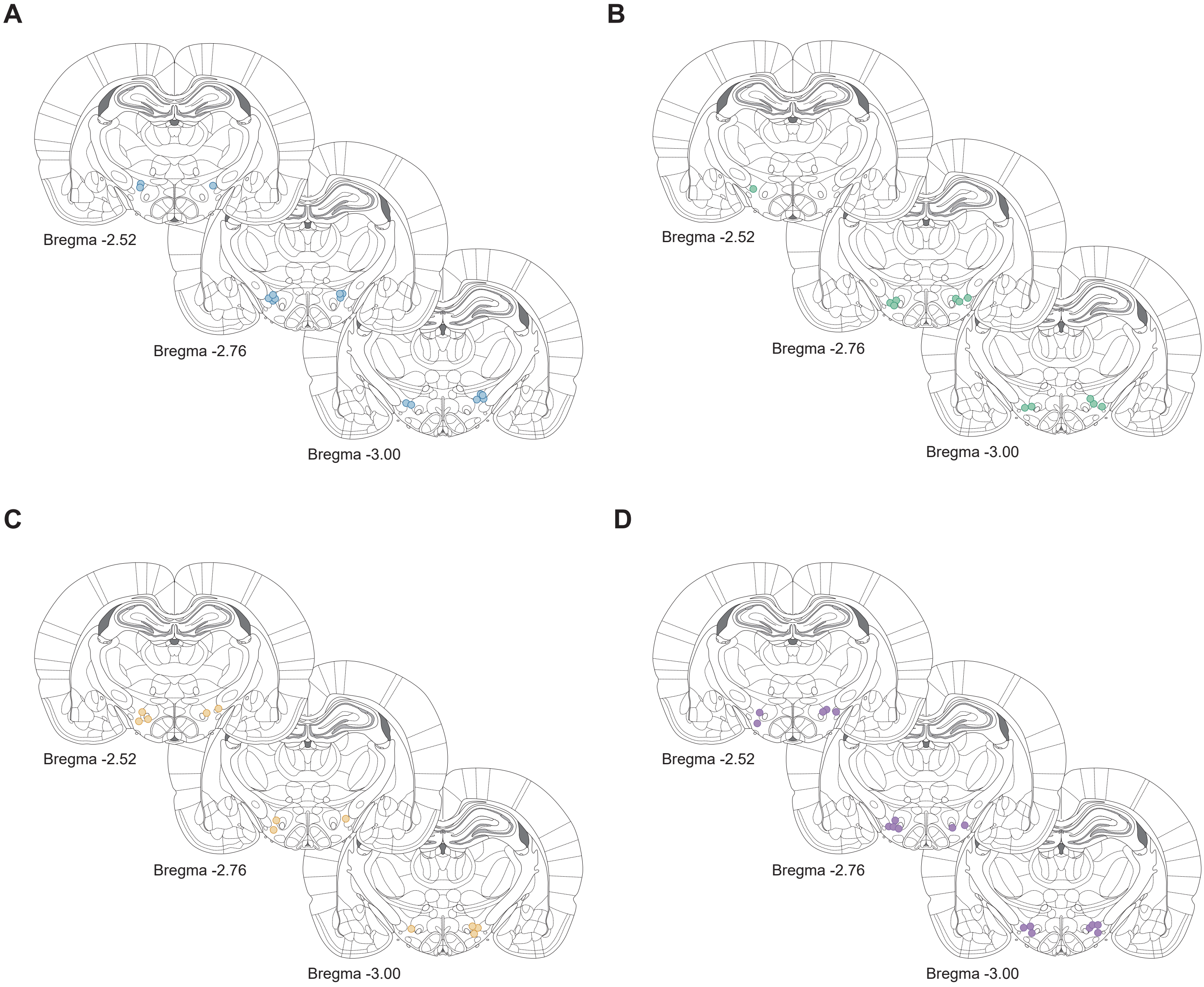
